# Tongue thickness measured by ultrasonography is associated with tongue pressure in the Japanese elderly

**DOI:** 10.1101/2020.02.26.966226

**Authors:** Masahiro Nakamori, Eiji Imamura, Masako Fukuta, Keisuke Tachiyama, Teppei Kamimura, Yuki Hayashi, Hayato Matsushima, Kanami Ogawa, Masami Nishino, Akiko Hirata, Tatsuya Mizoue, Shinichi Wakabayashi

## Abstract

The term “oral frailty” reflects the fact that oral health is associated with physical frailty and mortality. Tongue thickness measurement is one of the evaluations for swallowing function. The aim of this study was to investigate the relationship between tongue thickness and tongue pressure in the Japanese elderly. We evaluated 254 elderly patients; all the participants underwent tongue ultrasonography and tongue pressure measurement. To determine tongue thickness, we measured the vertical distance from the surface of the mylohyoid muscle to the tongue dorsum using ultrasonography. Result analyses revealed that tongue thickness was linearly associated with tongue pressure in both sexes. In male participants, dyslipidemia, lower leg circumference, and tongue pressure were independently significantly associated with tongue thickness. In female participants, body mass index and tongue pressure were independently significantly associated with tongue thickness. Tongue thickness was significantly decreased by age for both men and women. The optimal cutoff for tongue thickness to predict the tongue pressure of < 20 kPa was 41.3 mm in males, and 39.3 mm in females. In the Japanese elderly, tongue thickness using ultrasonography is associated with tongue pressure. Tongue thickness and tongue pressure, which are sensitive markers for oral frailty, decrease in aging. The tongue ultrasonography provides a less invasive technique for determining tongue thickness and related aspiration risks for elderly patients.

## Introduction

In an aging society, such as present-day Japan, frailty is a critical issue related to morbidity as well as mortality. Frailty is a common geriatric syndrome that embodies an elevated risk of catastrophic declines in health and function in the elderly. Frailty relates to many factors in an individual’s life, particularly physical, psychological, and social factors. Recently, it has been reported that oral health is associated with physical frailty and mortality, hence the term “oral frailty” [1-3]. Oral frailty consists of factors such as the diminished condition of teeth and ability to chew, swallow, and converse. It can lead to dysphagia, dehydration, malnutrition, asphyxia, and aspiration pneumonia, which is one of the most life-threatening concerns for the elderly [4,5].

The tongue is one of the most important organs related to oral frailty, a condition that can be life-threatening; swallowing dysfunction can lead to aspiration pneumonia and is a critical concern. Dysphagia is reported to be associated with oral, physical, cognitive, and psychological frailty in the elderly [6] and evaluations of swallowing function are important for assessing a person’s risk of aspiration or suffocation. Such evaluations may help determine decisions around eating and feeding, such as avoiding foods with certain textures to prevent aspiration pneumonia or suffocation and choosing foods with appropriate textures that can improve someone’s nutritional state. Quantitative evaluation of oral frailty often requires specialized instruments. Among them, videofluoroscopic examination (VF) is one of the most reliable and is seen as the gold standard for evaluating swallowing function. However, VF is a specialized method that is not routinely performed with the general population. Hence, several simple and non-invasive techniques have been developed for evaluating swallowing function, such as those for measuring tongue pressure and tongue thickness [7-9]. Tongue pressure is measured by having a patient raise up the tongue and push on the hard palate. Studies have reported that lower tongue pressure is a sensitive indicator for detecting swallowing dysfunction in patients who have had strokes and those with certain neurological disorders [10-12]. Of note, tongue pressure decreases with age and has been shown to be significantly lower in frail Japanese elderly persons [8].

Tongue thickness is often measured by ultrasonography, which can generate a stable numerical value [13,14]. It is reported that lower tongue pressure and decreasing tongue thickness measured by ultrasonography are associated with the oral preparatory and transit time measured using VF in amyotrophic lateral sclerosis patients [11,13]. Tongue ultrasonography is an objective and non-invasive evaluation technique that carries no risk of aspiration.

There are several reports of tongue pressure in the general population and the elderly, but not about how tongue pressure relates to tongue thickness measured by ultrasonography. The aim of this study was to investigate the relationship between tongue thickness and tongue pressure in the Japanese elderly.

## Methods

### Ethics Statement

The study protocols were approved by the ethics committee of Suiseikai Kajikawa Hospital and performed according to the guidelines of the national government based on the Helsinki Declaration of 1964. Written informed consent was obtained from all patients. All data analyses were blinded so no identifying information was revealed.

### Subjects

Consecutive outpatients who visited Suiseikai Kajikawa Hospital between February 1st and July 31st of 2019 were enrolled in this prospective study. We included those patients who consented to participate and were aged 65 years and older with chronic disease. We excluded patients who had a history of otorhinolaryngologic disease and/or neurodegenerative disease. Patients with paralysis were also excluded.

### Tongue ultrasonography

The non-invasive ultrasound examinations were performed by a neurosonologist (M Nakamori) using a Noblus imaging system (Hitachi, Ltd., Tokyo, Japan). Tongue thickness was measured using a 2-8MHz convex array transducer according to a previously reported method, with the device placed under the chin of the participant. The subjects were examined in a 30° reclining position while seated. Tongue thickness was determined by measuring the distance between the upper and lower surfaces of the patient’s lingual muscles in the center of the plane perpendicular to the Frankfurt horizontal plane of the frontal section (Fig 1A). This perpendicular plane intersects the distal surfaces of the bilateral mandibular second premolars. The vertical distance was measured from the surface of the mylohyoid muscle to the tongue dorsum (Fig 1B). This measurement was performed three times, and the mean value was defined as the tongue thickness for each participant. We confirmed the reliability of tongue ultrasonography by calculating the intra-rater and inter-rater reliability. For investigating intra-rater reliability, we measured the tongue thickness of the normal subject three times per day; their mean value was defined as the tongue thickness for the day. These measurements were repeated for ten days, and the resulting coefficient of variation was 1.59%. To investigate the inter-rater variability, two investigators (M Nakamori and MF) independently measured the tongue thickness of the same 23 normal subjects, and the resulting coefficient of variation was under 1.74%. Bland-Altman analysis was performed, and systematic (fixed and proportional) errors were not detected.

**Fig 1.**
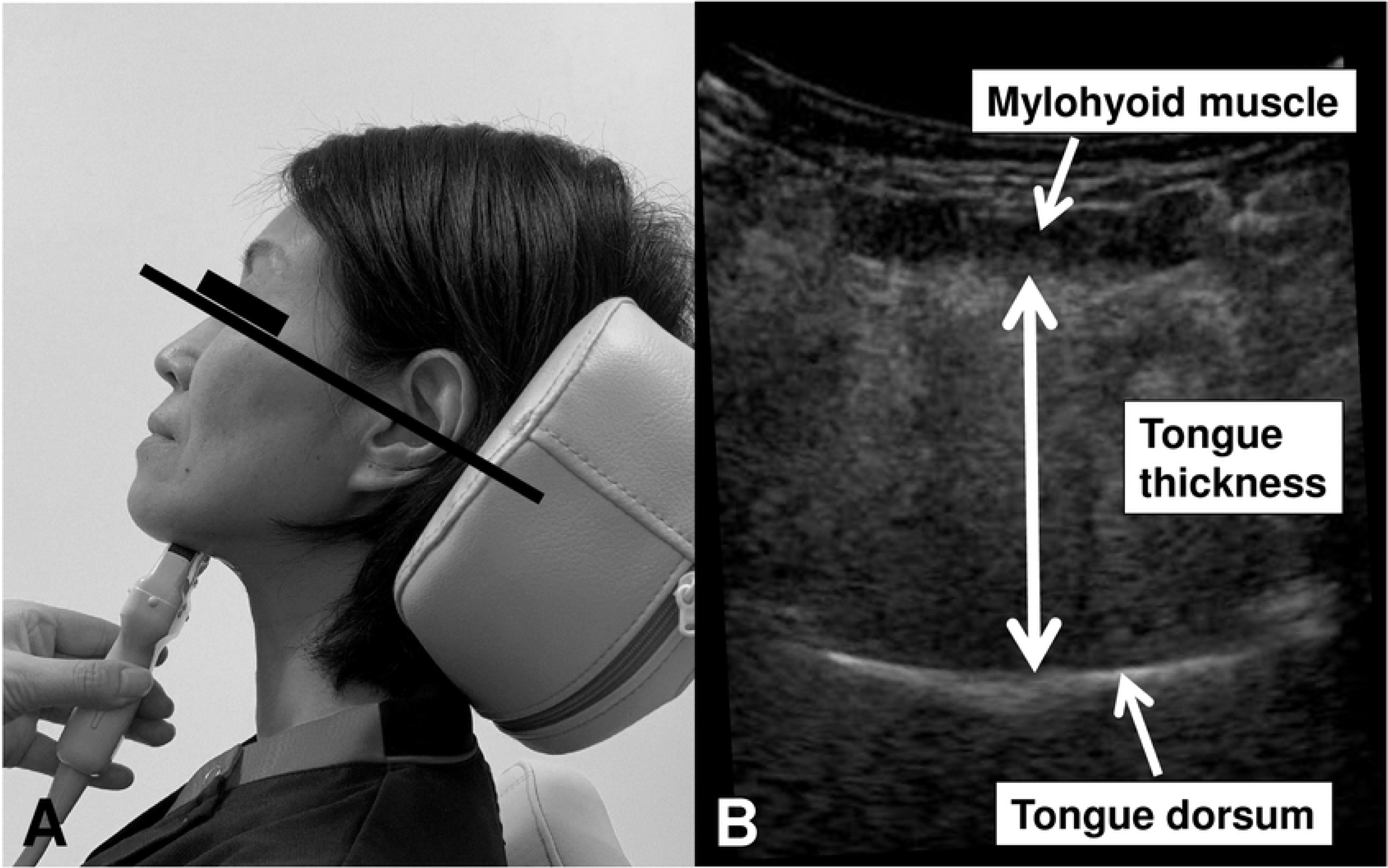
Measurement of tongue thickness. A) The subjects were examined in a 30° reclined position while seated. Tongue thickness was determined as the distance between the upper and lower surfaces of the lingual muscles in the center of the plane perpendicular to the Frankfurt horizontal plane in the frontal section. B) The vertical distance was measured from the surface of the mylohyoid muscle to the tongue dorsum.

### Tongue pressure measurement

Clinical technicians (KO, M Nishino, and AH) measured tongue pressure independently using balloon-type equipment (TPM-01; JMS Co. Ltd., Hiroshima, Japan) on the same day when the patients underwent measurement of tongue thickness using ultrasonography. The balloon-type equipment consisted of a disposable oral probe, an infusion tube as a connector, and a recording device (Fig 2A). For tongue pressure measurement, the subjects were placed in a relaxed sitting position and asked to place the balloon in their mouths, holding the plastic pipe at the midpoint of their central incisors with closed lips. The subjects were asked to maintain this position as clinicians adjusted the probe and confirmed that it was in the correct position. The subjects were then asked to raise their tongue and compress the small balloon with their palate at maximum voluntary effort for seven seconds as described previously (Fig 2B) [7,15]. This measurement was performed three times with the subjects resting for approximately 30 seconds and rinsing their mouths between each measurement. The highest value from the three measurements was defined as the tongue pressure for each subject. The reliability of intraindividual measurement was previously reported [10,16].

**Figure 2.**
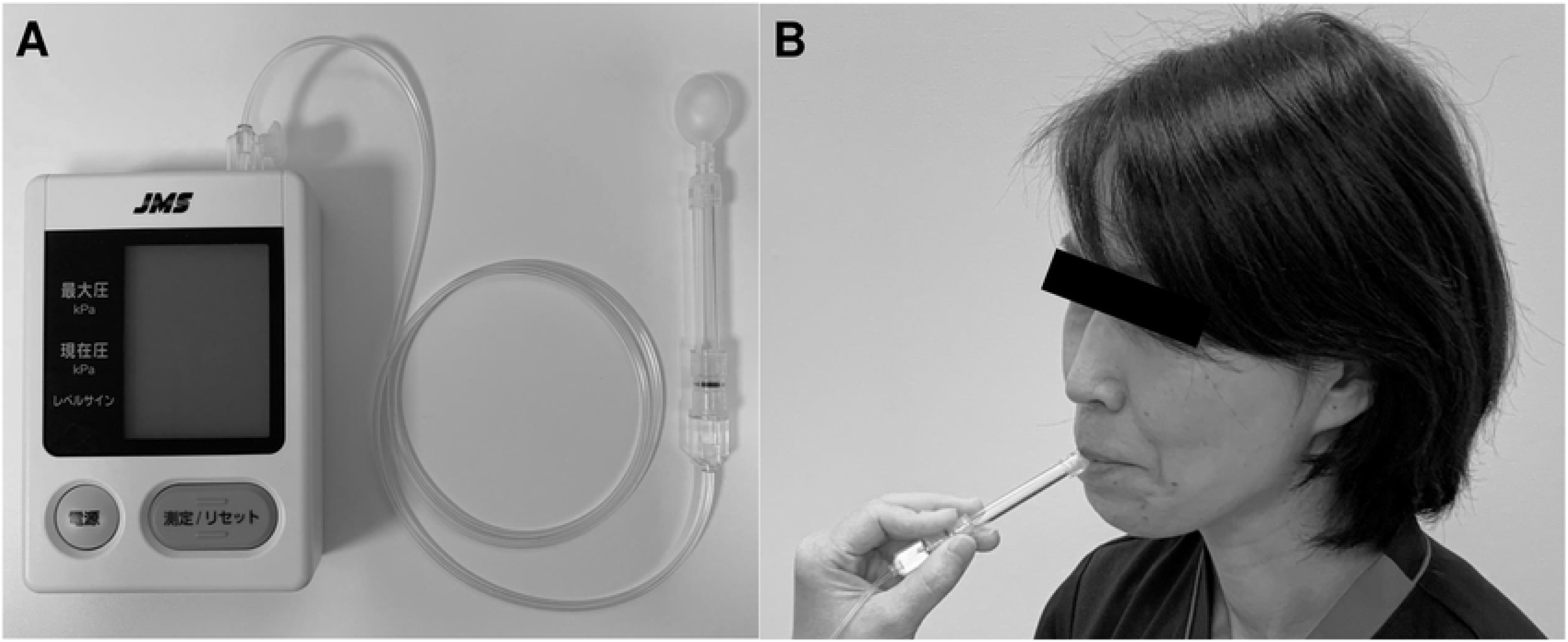
Measurement of tongue pressure. A) The balloon-type equipment consists of a disposable oral probe, an infusion tube as a connector, and a recording device. B) The subjects were asked to place the balloon in their mouths, holding the plastic pipe at the midpoint of their central incisors with closed lips. The subjects were asked to raise their tongue and compress the small balloon with their palate at maximum voluntary effort for 7 seconds.

### Data Acquisition

Patient characteristics, including age, gender, body mass index (BMI), past history of comorbidities (hypertension, diabetes mellitus, dyslipidemia, chronic kidney disease), grip power, lower leg circumference, serum albumin, tongue pressure, and tongue thickness were evaluated. Hypertension was defined as the use of anti-hypertensive medication or confirmed blood pressure of ≥ 140/90 mmHg at rest. Diabetes mellitus was defined as a glycated hemoglobin level of ≥ 6.5%, fasting blood glucose level of ≥ 126 mg/dl, or use of anti-diabetes medication. Dyslipidemia was defined as a total cholesterol level of ≥ 220 mg/dl, low-density lipoprotein cholesterol level of ≥ 140 mg/dl, high-density lipoprotein cholesterol level of < 40 mg/dl, triglyceride levels of ≥ 150 mg/dl, or use of anti-hyperlipidemia medication. Renal functioning was calculated with the estimated glomerular filtration rate (eGFR) using a revised equation for the Japanese population as follows: eGFR (ml min^−1^ 1.73 m^−2^) = 194 × (serum creatinine)^−1.094^ × (age)^−0.287^ × 0.739 (for women) [17]. Chronic kidney disease was defined as an eGFR < 60 ml min^−1^ 1.73 m^−2^. Grip power was measured for both sides and the mean value was used for analysis. Lower leg circumference was measured at the thickest place at both sides and the mean value was used for analysis.

### Statistical analysis

The data were expressed as the mean ± standard deviation for continuous variables and frequencies and percentages for discrete variables. Statistical analysis was performed using JMP 13 statistical software (SAS Institute Inc., Cary, NC, USA). The statistical significance of intergroup differences was assessed using unpaired *t-*tests or χ^2^ tests as appropriate. The baseline data in the subjects were analyzed, and two-step strategies were employed to assess the relative importance of variables in their association with tongue thickness using least square linear regression analysis. First, a univariate analysis was performed. Then, a multifactorial least-square linear regression analysis was performed with selected factors that had *p* < 0.20 on univariate analysis. Tongue thickness and tongue pressure were compared by five-year age increments. The data were analyzed with a one-way analysis of variance and Tukey’s honestly significant difference (HSD) test. Receiver operating characteristic (ROC) analysis was performed to determine the tongue thickness predicting a tongue pressure < 20kPa, which suggests swallowing dysfunction. We considered *p* < 0.05 as statistically significant.

## Results

We evaluated 254 elderly patients, whose backgrounds are shown in Table 1. Tongue thickness, tongue pressure, grip power, and lower leg circumference were all markedly different between the male group and the female group. To account for a disproportionate physique owing to normal differences, between males and females, results were analyzed separately by sex.

**Table 1.** Japanese Elder Participants’ Health-Related Factors in Tongue Thickness Study.

|  | All<br>N = 254 | Male<br>N = 163 | Female<br>N = 91 | *p*-value |
| --- | --- | --- | --- | --- |
| **Age** | 77.9 ±6.3 | 76.9 ±5.8 | 79.5 ±6.7 | 0.001 |
| **Body mass index, kg/m<sup>2</sup>** | 23.2 ±3.0 | 23.3 ±2.8 | 23.2 ±3.2 | 0.776 |
| **Hypertension, n (%)** | 207 (81.5) | 129 (79.1) | 78 (85.7) | 0.196 |
| **Diabetes mellitus, n (%)** | 51 (20.0) | 35 (21.5) | 16 (17.6) | 0.458 |
| **Dyslipidemia, n (%)** | 172 (67.7) | 104 (63.8) | 68 (74.7) | 0.074 |
| **Chronic kidney disease, n (%)** | 48 (18.9) | 33 (20.3) | 15 (16.5) | 0.463 |
| **grip power, kg** | 24.2 ±8.7 | 28.8 ±6.9 | 15.9 ±4.5 | < 0.001 |
| **lower leg circumference, cm** | 33.6 ±3.3 | 34.4 ±3.2 | 32.1 ±2.8 | < 0.001 |
| **Serum albumin, g/dl** | 4.1 ±0.3 | 4.1 ±0.3 | 4.1 ±0.4 | 0.345 |
| **Tongue pressure, kpa** | 35.7 ±10.6 | 37.4 ±10.2 | 32.5 ±10.5 | < 0.001 |
| **Tongue thickness, mm** | 41.9 ±2.8 | 42.4 ±2.6 | 40.9 ±2.8 | < 0.001 |

Scatter plots were used to indicate tongue thickness and tongue pressure by sex. These analyses indicate that tongue thickness was linearly associated with tongue pressure in both sexes (male; coefficient 0.202, 95% confidence interval 0.182–0.223, *p* < 0.001; Fig 3A, female; coefficient 0.202, 95% confidence interval 0.182–0.223, *p* < 0.001; Fig 3B).

**Fig 3.**
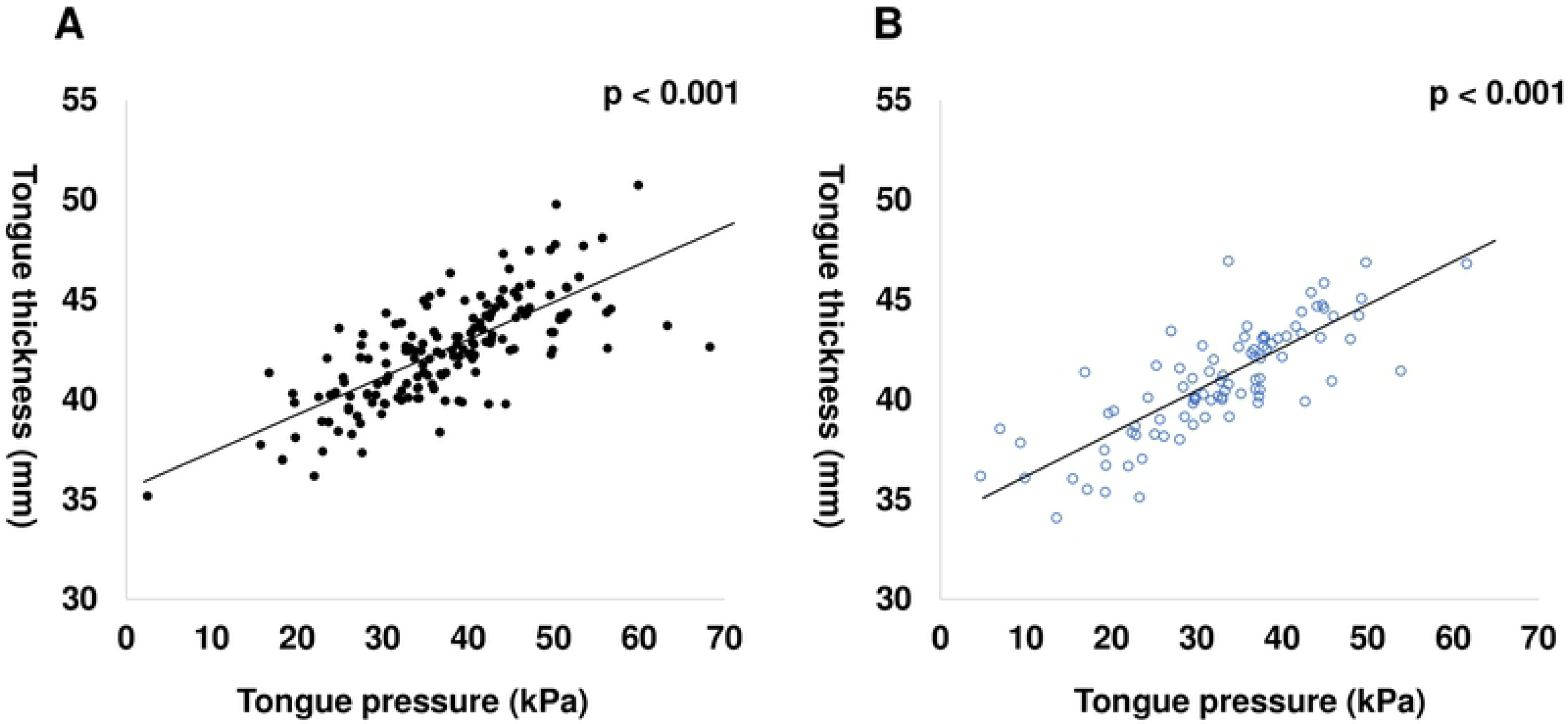
The Association of Tongue Thickness with Tongue Pressure. In the elderly, tongue thickness is shown to be associated with tongue pressure A) in males (*p* < 0.001) and B) in females (*p* < 0.001); ●, male; ○, female.

The potential factors associated with tongue thickness (listed in Table 1) were evaluated using multifactorial regression analysis by sex. In the male group, dyslipidemia, lower leg circumference, and tongue pressure were independently significant in their association with tongue thickness (adjusted R^2^ = 0.653, *p* < 0.001, n = 163) (Table 2A). In the female group, body mass index and tongue pressure were independently significant in their association with tongue thickness (adjusted R^2^ = 0.707, *p* < 0.001, n = 91) (Table 2B).

**Table 2.** Factors Influencing Tongue Thickness in Japanese Elders.

TONGUE THICKNESS AND TONGUE PRESSURE IN THE ELDERLY
| Male | Univariate | Multivariate |  |  |
| --- | --- | --- | --- | --- |
|  | *p*-value | coefficient | 95% CI | *p*-value |
| Age | < 0.001 | -0.001 | -0.050 - 0.047 | 0.962 |
| Body mass index, kg/m<sup>2</sup> | < 0.001 | 0.070 | -0.057 - 0.198 | 0.277 |
| Hypertension, n (%) | 0.937 |  |  |  |
| Diabetes mellitus, n (%) | 0.174 | -0.266 | -0.564 - 0.032 | 0.080 |
| Dyslipidemia, n (%) | 0.032 | -0.264 | -0.519 - -0.001 | 0.043 |
| Chronic kidney disease, n (%) | 0.406 |  |  |  |
| grip power, kg | < 0.001 | 0.032 | -0.011 - 0.075 | 0.141 |
| lower leg circumference, cm | < 0.001 | 0.192 | 0.065 - 0.319 | 0.003 |
| Serum albumin, g/dl | < 0.001 | 0.541 | -0.391 - 1.474 | 0.253 |
| Tongue pressure, kPa | < 0.001 | 0.142 | 0.114 - 0.169 | < 0.001 |

B
| Female | Univariate | Multivariate |  |  |
| --- | --- | --- | --- | --- |
|  | *p*-value | coefficient | 95% CI | *p*-value |
| Age | < 0.001 | -0.004 | -0.067 – 0.059 | 0.896 |
| Body mass index, kg/m<sup>2</sup> | < 0.001 | 0.195 | 0.053 - 0.337 | 0.008 |
| Hypertension, n (%) | 0.776 |  |  |  |
| Diabetes mellitus, n (%) | 0.336 |  |  |  |
| Dyslipidemia, n (%) | 0.989 |  |  |  |
| Chronic kidney disease, n (%) | 0.474 |  |  |  |
| grip power, kg | < 0.001 | 0.073 | -0.019 – 0.164 | 0.118 |
| lower leg circumference, cm | < 0.001 | 0.082 | -0.094 – 0.258 | 0.359 |
| Serum albumin, g/dl | 0.077 | 0.394 | -0.569 – 1.357 | 0.419 |
| Tongue pressure, kPa | < 0.001 | 0.171 | 0.135–0.207 | < 0.001 |

Tongue thickness was compared by five-year age increments for each sex (Fig 4). These data suggested that tongue thickness is significantly decreased by age in each sex (*p* < 0.001). In addition, tongue pressure was compared by five-year age increments for each sex (Fig 5). These data suggested that tongue pressure also significantly decreased by age in each sex (*p* < 0.001). Tukey’s HSD tests revealed that both tongue thickness and tongue pressure remarkably decreased in patients over 85 years old.

**Fig 4.**
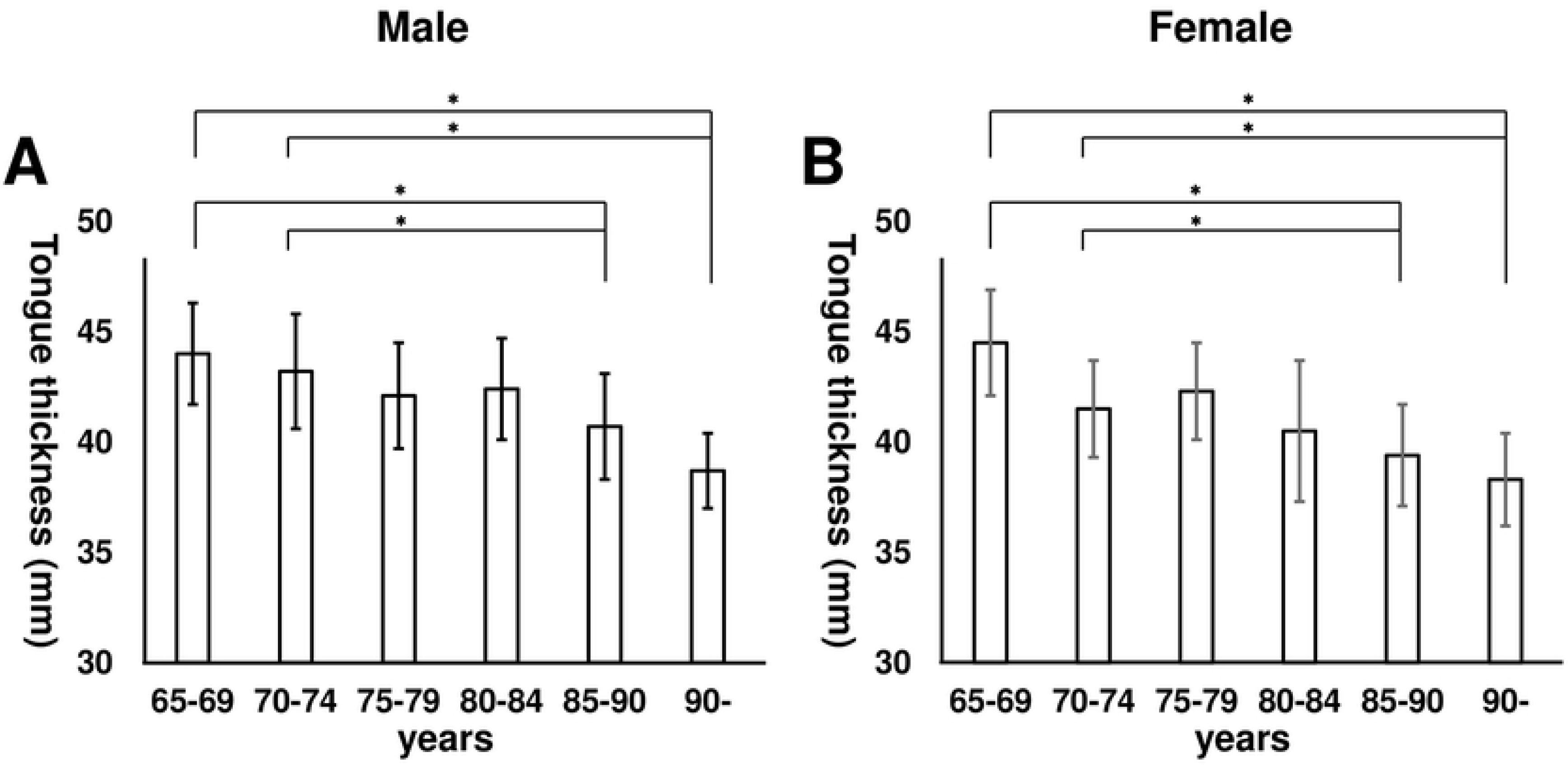
Tongue Thickness by Age and Sex. A) Tongue thickness by five-year age increments in males. Tongue thickness significantly decreased with increased age (*p* < 0.001). B) Tongue thickness by five-year age increments in females. Tongue thickness was significantly decreased with increased age (*p* < 0.001). \**p* < 0.05 by Tukey’s honestly significant difference test.

**Fig 5.**
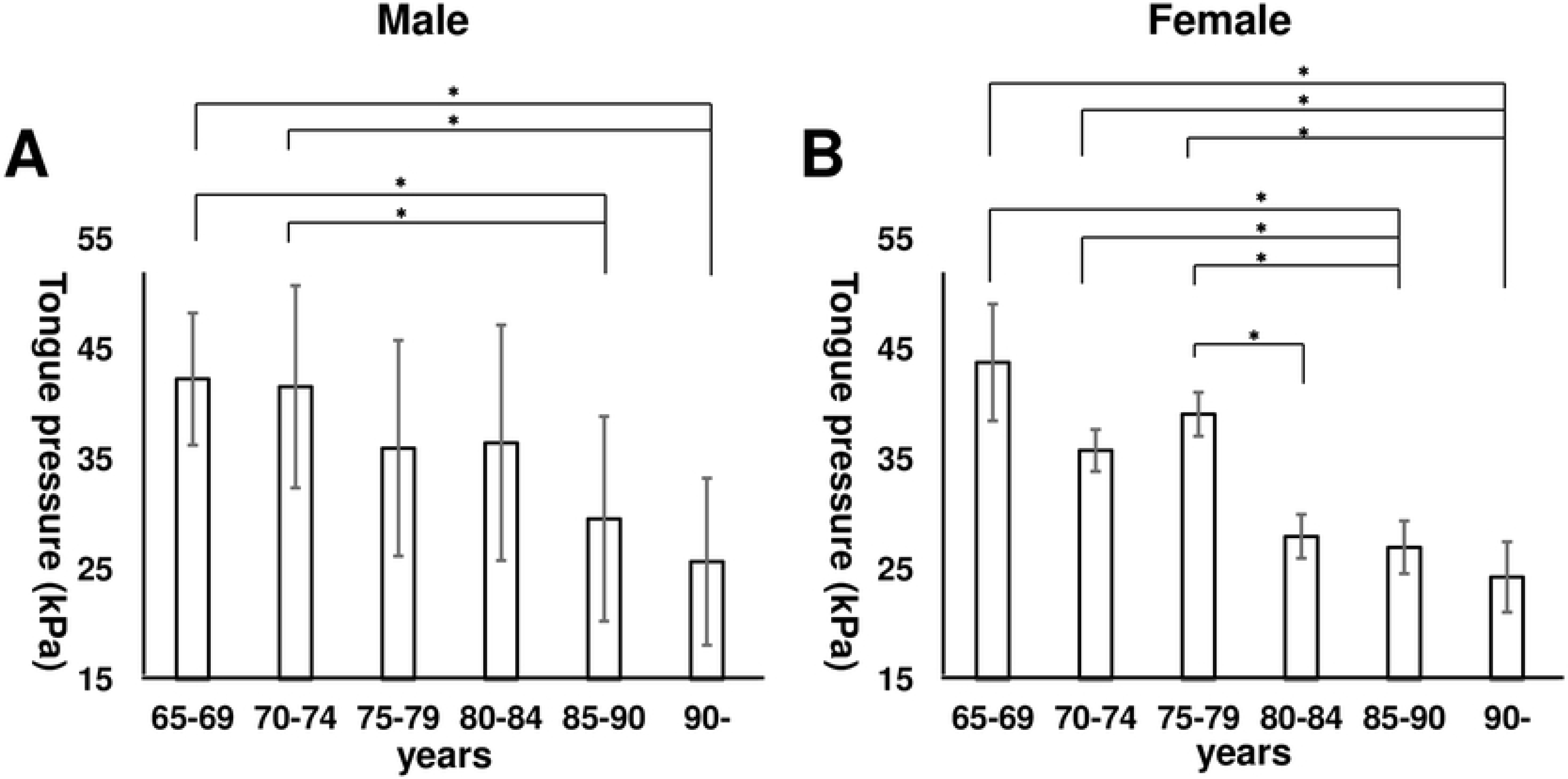
Tongue Pressure by Age and Sex. A) Tongue pressure by five-year age increments in males. Tongue pressure significantly decreased with increased age (*p* < 0.001). B) Tongue pressure by five-year age increments in females. Tongue pressure significantly decreased with increased age (*p* < 0.001). \**p* < 0.05 by Tukey’s honestly significant difference test.

Previous several reports suggested that using the JMS balloon-type device, the tongue pressure of patients with swallowing dysfunction was observed to be approximately < 20 kPa [10-12]. The optimal cutoff for tongue thickness to predict the tongue pressure < 20 kPa was 41.3 mm in male from the ROC analysis (χ^2^ = 24.48, *p* < 0.001, sensitivity 100.0%, specificity 68.4%, AUC = 0.91), and 39.3 mm in female from the ROC analysis (χ^2^ = 32.29, *p* < 0.001, sensitivity 91.7%, specificity 82.3%, AUC = 0.92).

## Discussion

In the present study, we investigated the tongue thickness of elder Japanese patients using ultrasonography. Measurements of tongue thickness and tongue pressure are useful evaluations for detecting swallowing function in patients with neurological diseases [10-13]. Prior studies were conducted to examine tongue pressure measured in the general population, revealing that tongue pressure gradually decreases with age, and markedly decreases in frail elderly people [8,18]. In addition, decreased tongue pressure is associated with sarcopenia and sarcopenic dysphagia in the elderly [19]. On the other hand, tongue thickness has been scarcely evaluated in the general population. We evaluated tongue thickness using ultrasonography in reasonably healthy elderly patients with no neurological deficits.

In this study, we focused on reasonably healthy elderly patients who have a few lifestyle diseases. Our results showed that, for the elderly participants, tongue thickness was independently associated with tongue pressure and lower leg circumference in males, and tongue pressure and body mass index in females. A decrease of lower leg circumference or body mass index is also known to be related to sarcopenia which leads to frailty. In addition, it has been reported that tongue pressure is significantly reduced in frail elderly Japanese persons [8]. Hence, decreased tongue thickness may also be one of the signs of frailty.

Tongue thickness and tongue pressure significantly decreased with age in each sex. in our analysis, the decrease was most remarkable in participants over 85 years old. These results allude to the rate of progression of tongue muscle atrophy due to aging. We generally judge tongue atrophy by visual examination, but it is fairly subjective, depending on the opinion of the examiner. In contrast, measurement with ultrasonography is quantitative and objective. Several previous reports have suggested that, using the JMS balloon-type device, the tongue pressure of patients with swallowing dysfunction was observed to be approximately < 20 kPa [10-12]. ROC analysis of the results of our study revealed that the optimal cutoff for tongue thickness to predict tongue pressure of < 20 kPa is 41.3 mm for males and 39.3 mm for females.

Oral frailty consists of many factors such as the condition of teeth and the ability to chew, swallow, converse, and so on. For all these factors, the tongue plays a major role and thereby tongue condition is a key component of oral frailty. One serious concern related to oral frailty is that elderly people often developed aspiration pneumonia. In Japan, aspiration pneumonia is a major cause of death and its risk is increased in the elderly [20]. While tongue pressure as measured at approximately 20 kPa is a cutoff value for swallowing dysfunction, many people who meet that description do not feel significantly orally dysfunctional or experience dysphagia [11,12]. For early detection, measurements of tongue thickness and tongue pressure should be performed as routine screening measures in the general elderly population. However, the measurement of tongue pressure requires a patient to be capable of comprehending the process and to actively participate while following instructions. Patients with dementia, for example, may not be able to undergo tongue pressure measurements. For these patients, tongue thickness measurement might be useful as a more suitable screening test. Early detection of tongue hypofunction can prevent the progression of oral frailty— which can lead to systemic frailty—and can improve frailty-related morbidity and mortality. Rehabilitation for raising tongue pressure has been developed and reported as effective for the general elderly population [21]. Moreover, it has been shown that the maintenance of body weight by nutritional intervention may inhibit the progression of tongue atrophy [13]. Nutritional care is also important for preventing oral and systemic frailty.

This study, while encouraging, also has limitations. First, the measurements were not compared with actual oral dysfunction or gold standard methods such as VF. In amyotrophic lateral sclerosis patients, it has been reported that tongue thickness and tongue pressure show an association in VF temporary analysis, especially oral preparatory and transit time [11,13]. However, in the general elderly, there are only limited opportunities to perform VF. Second, in the present study, tongue thickness of older participants was not compared with that of younger healthy subjects. Data exists regarding tongue pressure, as it has been measured in all ages. Further studies are needed to measure the tongue thickness in a general population including all ages.

## Conclusion

In the Japanese elderly, tongue thickness, as measured using ultrasonography, is associated with tongue pressure. Decreases in both tongue thickness and tongue pressure occur in aging patients and may be sensitive markers of oral frailty. Oral frailty decreased tongue pressure, and decreased tongue thickness lead to systemic frailty and, finally, affect morbidity and mortality. Early detection using such instruments might be important in preventing the progression of frailty.

## Acknowledgments

We would like to sincerely thank the staff at the Suiseikai Kajikawa Hospital for their technical assistance.

## References

1. Rapp L, Sourdet S, Vellas B, Lacoste-Ferre MH. Oral Health and the Frail Elderly. J Frailty Aging. 2017;6: 154–160. doi: 10.14283/jfa.2017.9.

2. Tanaka T, Takahashi K, Hirano H, Kikutani T, Watanabe Y, Chara Y, et al. Oral Frailty as a Risk Factor for Physical Frailty and Mortality in Community-Dwelling Elderly. J Gerontol A Biol Sci Med Sci. 2018;73: 1661–1667. doi: 10.1093/gerona/glx225.

3. Hakeem FF, Bernabe E, Sabbah W. Association between oral health and frailty: A systematic review of longitudinal studies. Gerodontology. 2019;36: 205–215. doi: 10.1111/ger.12406.

4. Hudson HM, Daubert CR, Mills RH. The interdependency of protein-energy malnutrition, aging, and dysphagia. Dysphagia. 2000;15: 31–38. DOI: 10.1007/s004559910007.

5. Marik PE, Kaplan D. Aspiration pneumonia and dysphagia in the elderly. Chest. 2003;124: 328–336. DOI: 10.1378/chest.124.1.328.

6. Nishida T, Yamabe K, Honda S. Dysphagia is associated with oral, physical, cognitive and psychological frailty in Japanese community-dwelling elderly persons. Gerodontology. 2019; doi: 10.1111/ger.12455.

7. Yoshikawa M, Yoshida M, Tsuga K, Akagawa Y, Groher ME. Comparison of three types of tongue pressure measurement devices. Dysphagia. 2011;26: 232–237. doi: 10.1007/s00455-010-9291-3.

8. Tsuga K, Yoshikawa M, Oue H, Okazaki Y, Tsuchioka H, Maruyama M, et al. Maximal voluntary tongue pressure is decreased in Japanese frail elderly persons. Gerodontology. 2012;29: e1078–1085.

9. Weikamp JG, Schelhaas HJ, Hendriks JC, de Swart BJ, Geurts AC. Prognostic value of decreased tongue strength on survival time in patients with amyotrophic lateral sclerosis. J Neurol. 2012;259: 2360–2365. doi: 10.1007/s00415-012-6503-9.

10. Nakamori M, Hosomi N, Ishikawa K, Imamura E, Shishido T, Ohshita T, et al. Prediction of Pneumonia in Acute Stroke Patients Using Tongue Pressure Measurements. PLoS One. 2016;11: e0165837. doi: 10.1371/journal.pone.0165837.

11. Hiraoka A, Yoshikawa M, Nakamori M, Hosomi N, Nagasaki T, Mori T, et al. Maximum Tongue Pressure is Associated with Swallowing Dysfunction in ALS Patients. Dysphagia. 2017;32: 542–547. doi: 10.1007/s00455-017-9797-z.

12. Mano T, Katsuno M, Banno H, Suzuki K, Suga N, Hashizume A, et al. Tongue pressure as a novel biomarker of spinal and bulbar muscular atrophy. Neurology. 2014;82: 255–262. doi: 10.1212/WNL.0000000000000041.

13. Nakamori M, Hosomi N, Takaki S, Oda M, Hiraoka A, Yoshikawa, M, et al. Tongue thickness evaluation using ultrasonography can predict swallowing function in amyotrophic lateral sclerosis patients. Clin Neurophysiol. 2016;127: 1669–1674. doi: 10.1016/j.clinph.2015.07.032.

14. Tamura F, Kikutani T, Tohara T, Yoshida M, Yaegaki K. Tongue thickness relates to nutritional status in the elderly. Dysphagia. 2012;27: 556–561. doi: 10.1007/s00455-012-9407-z.

15. Tsuga K, Maruyama M, Yoshikawa M, Yoshida M, Akagawa Y. Manometric evaluation of oral function with a hand-held balloon probe. J Oral Rehabil. 2011;38: 680–685. doi: 10.1111/j.1365-2842.2011.02202.x.

16. Hayashi R, Tsuga K, Hosokawa R, Yoshida M, Sato Y, Akagawa Y. A novel handy probe for tongue pressure measurement. Int J Prosthodont. 2002;15: 385–388.

17. Matsuo S, Imai E, Horio M, Yasuda Y, Tomita K, Nitta K, et al. Revised equations for estimated GFR from serum creatinine in Japan. Am J Kidney Dis. 2009;53: 982–992. doi: 10.1053/j.ajkd.2008.12.034.

18. Utanohara Y, Hayashi R, Yoshikawa M, Yoshida M, Tsuga K, Akagawa Y. Standard values of maximum tongue pressure taken using newly developed disposable tongue pressure measurement device. Dysphagia. 2008;23: 286–290. doi: 10.1007/s00455-007-9142-z.

19. Maeda K, Akagi J. Decreased tongue pressure is associated with sarcopenia and sarcopenic dysphagia in the elderly. Dysphagia. 2015;30: 80–87. doi: 10.1007/s00455-014-9577-y.

20. Teramoto S, Fukuchi Y, Sasaki H, Sato K, Sekizawa K, Matsuse T, et al. High incidence of aspiration pneumonia in community- and hospital-acquired pneumonia in hospitalized patients: a multicenter, prospective study in Japan. J Am Geriatr Soc. 2008;56: 577–579. doi: 10.1111/j.1532-5415.2008.01597.x.

21. Van den Steen L, De Bodt M, Guns C, Elen R, Vanderwegen J, Van Nuffelen G. Tongue-Strengthening Exercises in Healthy Older Adults: Effect of Exercise Frequency - A Randomized Trial. Folia Phoniatr Logop. 2020; doi: 10.1159/000505153.

